# Crowdsourcing RNA structural alignments with an online computer game

**DOI:** 10.1101/009902

**Authors:** Jérôme Waldispühl, Arthur Kam, Paul P. Gardner

**Affiliations:** School of Computer Science, McGill University, Montreal, QC H3A 0E9, Canada; Biomolecular Interaction Centre, School of Biological Science, University of Canterbury, Christchurch, New Zealand

**Keywords:** Crowd-sourcing, Human-computing game, RNA alignment

## Abstract

The annotation and classification of ncRNAs is essential to decipher molecular mechanisms of gene regulation in normal and disease states. A database such as `Rfam` maintains alignments, consensus secondary structures, and corresponding annotations for RNA families. Its primary purpose is the automated, accurate annotation of non-coding RNAs in genomic sequences. However, the alignment of RNAs is computationally challenging, and the data stored in this database are often subject to improvements. Here, we design and evaluate `Ribo`, a human-computing game that aims to improve the accuracy of RNA alignments already stored in `Rfam`. We demonstrate the potential of our techniques and discuss the feasibility of large scale collaborative annotation and classification of RNA families.

## 1. Introduction

Non-coding RNAs (ncRNAs) are functional RNA molecules that are not translated into proteins. They play key roles in aspects of gene transcription, protein transport, molecular assembly and regulatory processes (e.g. riboswitches and microRNAs).^1,2^ The annotation and classification of ncRNAs is essential to decipher molecular mechanisms of gene regulation in normal and disease states. The `Rfam` database maintains alignments, consensus secondary structures, and corresponding annotations for RNA families. Its primary purpose is the automated, accurate annotation of non-coding RNAs in genomic sequences.^3^ However, the alignments stored in this database are often subject to improvements. In fact, the `Rfam` consortium recently released a open call for participation, asking to its users to submit new or improved RNA alignments (http://rfam.sanger.ac.uk/submit_alignment).

Initiatives such as OpenStreetMap and crowdcrafting have proven that crowd-sourcing and human-computing techniques are valuable ways to both analyze and annotate large datasets that require human expertise as well as to solve problems that are difficult to treat with classical computer algorithms. Scientific games like `Foldit`^4^ and our previous contribution `Phylo`^5^ illustrate the potential of these techniques for studying, mining, and processing molecular biology data. More recent applications of scientific games to molecular biology problems include `Dizeez`,^6^ `The Cure`,^7^ `EteRNA`,^8^ `Fraxinus`,^9^ and `Nanocrafter`.^10^ Currently, people collectively spend an estimated 3 billions hours per week playing computer games. By tapping into this tremendous source of human attention and effort, human-computing games have the potential to bring massive resources to bear on solving complex problems arising in genomics.^11^

This call emphasizes the potential impact of new tools such as `Phylo` in genomics. The number of new RNA discoveries has accelerated thanks to new sequencing technologies and computational tools,^12–18^ consequently the `Rfam` curators are now overwhelmed by the sheer number of ncRNAs that require their attention.^19^ This is largely driven by the discovery of thousands of ncRNAs using RNA-seq datasets.^18,20^ Functional validation is also carried out using high-throughput approaches such as transposon mutagenesis^15,21^ and large-scale genome project provides useful evolutionary conservation dimension.^22,23^ The small resources of the `Rfam` consortium are significantly stretched with keeping the database up to date with new families, revisiting existing families with new information and maintaining the core Rfam resources such as the website, MySQL database and the different data queries. Some of this has been aleviated by contributions from the community via the RNA Families track with the journal RNA Biology.^24,25^ However, additional inputs from the research community such as crowd-sourcing and human-computing techniques could be valuable to maintain the quality of the data.

One of the difficulties for RNA analysis is that RNA sequences are generally poorly conserved whereas RNA structures are generally conserved.^26^ The bulk of these structures are determined by secondary structure interactions.^27^ These are formed by hydrogen bonding interactions between nucleotides (A-U, C-G and G-U) and basepair stacking interactions. Consequently, the tools for aligning homologous RNAs need to take structure into account in order to make accurate predictions. However, this is an NP-complete optimization problem^28^ and to-date no ideal heuristic solution has been implemented.^29^ Therefore, the “gold standard” for RNA sequence analysis remains the manual refinement of RNA alignments which have produced highly accurate structure predictions. In extreme cases, 97-98% of manually inferred structures were validated by crystallographic methods.^30^

Our group was the first to bring citizen science to the field of comparative genomics when, in 2010, we released `Phylo` (http://phylo.cs.mcgill.ca), a human-computing framework to solve the multiple sequence alignment (MSA) problem. The key idea of `Phylo` is to convert the MSA problem into a casual puzzle game that can be played by ordinary web users with a minimal prior knowledge of the biological context. In our original study,^5^ the puzzles were extracted from a 44-species MSA stored at the UCSC genome browser, and the best solutions have been re-inserted at their original locations to produce a higher quality MSA. One of the main innovations of `Phylo` was to push the gamification aspect at its limits. Unlike `Foldit` that require each new player to learn the basics of the biophysics of protein folding through a detailed tutorial before starting to play, `Phylo` is a true casual game, requiring absolutely no knowledge of genomics. Indeed, the latter is an intuitive Tetris-like game where players have to match colored blocks. As a consequence, the game is accessible to a broader audience and can benefit of the workforce provided by crowds composed of ordinary web users with a minimal prior knowledge of the biological context.

Here, we design and experiment with `Ribo`, a human-computing game that aims to improve the accuracy of RNA alignments. ncRNAs are characterized by a conserved secondary structure associated with their function. Therefore, RNA alignments require to simultaneously align sequences and secondary structures. We propose to develop a game inspired from `Phylo` for this specific case. We introduce new types of blocks representing the base-pairs of the secondary structure. Our working prototype uses left- and right-handed triangles to represent open and closing base-pairs of the bracket notation. In addition, unlike `Phylo`, this game is not using a phylogenetic tree and thus is easier to understand for non-experienced players. We evaluate the quality of the alignments calculated by the players with the `Infernal` package.^31^ In particular, we show that the solutions we collected through `Ribo` enabled us to build covariance models with better overall homolog recognition performances than the ones built from the initial `Rfam` alignments. This work suggests that the use of human-computing games has the potential to become a valuable resource to maintain RNA alignment databases. `Ribo` is available at http://ribo.cs.mcgill.ca.

## 2. Methods

### 2.1. RNA secondary structures and RNA alignments

Ribonucleic acids (RNAs) are versatile biomolecules that are involved a diverse number of biological functions. For example, as messenger RNA it encodes genes, as microRNA it regulates genes and as ribosomal RNA it translates genes. To achieve their functions, non-coding RNAs (ncRNAs) use sophisticated structures that can be described at two levels. First, the secondary structure is the set of all canonical base-pairing interactions found in the native conformation of the molecule. The canonical base pairs include Watson-Crick interactions between Adenine (A) and Uracil (U) or Guanine (G) and Cytosine (C), as well as Wobble interactions between Guanine and Uracil bases. Contiguous canonical base pairs form secondary structure elements called helices (or stems) connected together through various types of loops (E.g. hairpins, bulges, internal loops and multi-loops). The majority of secondary structures can be represented at planar graphs. Moreover, less than 5% of the secondary structures found in the `Rfam` database^3^ contain crossing interactions also called pseudo-knots. Hence, many secondary structures can be conveniently represented using the dot-brack notation that is illustrated in Figure 1. Then, the secondary structure elements are assembled together via numerous van der Waals contacts and specific hydrogen bonds into the tertiary structure (or 3D structure). RNA folding is hierarchical. The secondary structures of RNA form rapidly, these act as a scaffold for the slower formation of tertiary structures.^27^ For this reason, secondary structures provide a relatively accurate signature of the molecular function, as illustrated by the strong evolutionary conservation of RNA secondary structures.^32^ For instance, the secondary structures of tRNAs adopt a typical cloverleaf shape.^33,34^ The evolution of homologous ncRNAs is constrained by these functional structures, and their alignments generally comply with this information too. Thus, a ncRNA alignment is associated with a consensus secondary structure that is representative of the functional family. To date, the `Rfam` database is the most popular repository of structured ncRNA alignments.^3^ These alignments can then used to build covariance models, which are widely used for the functional annotation of ncRNA sequences with unknown functions.

**Fig. 1.**
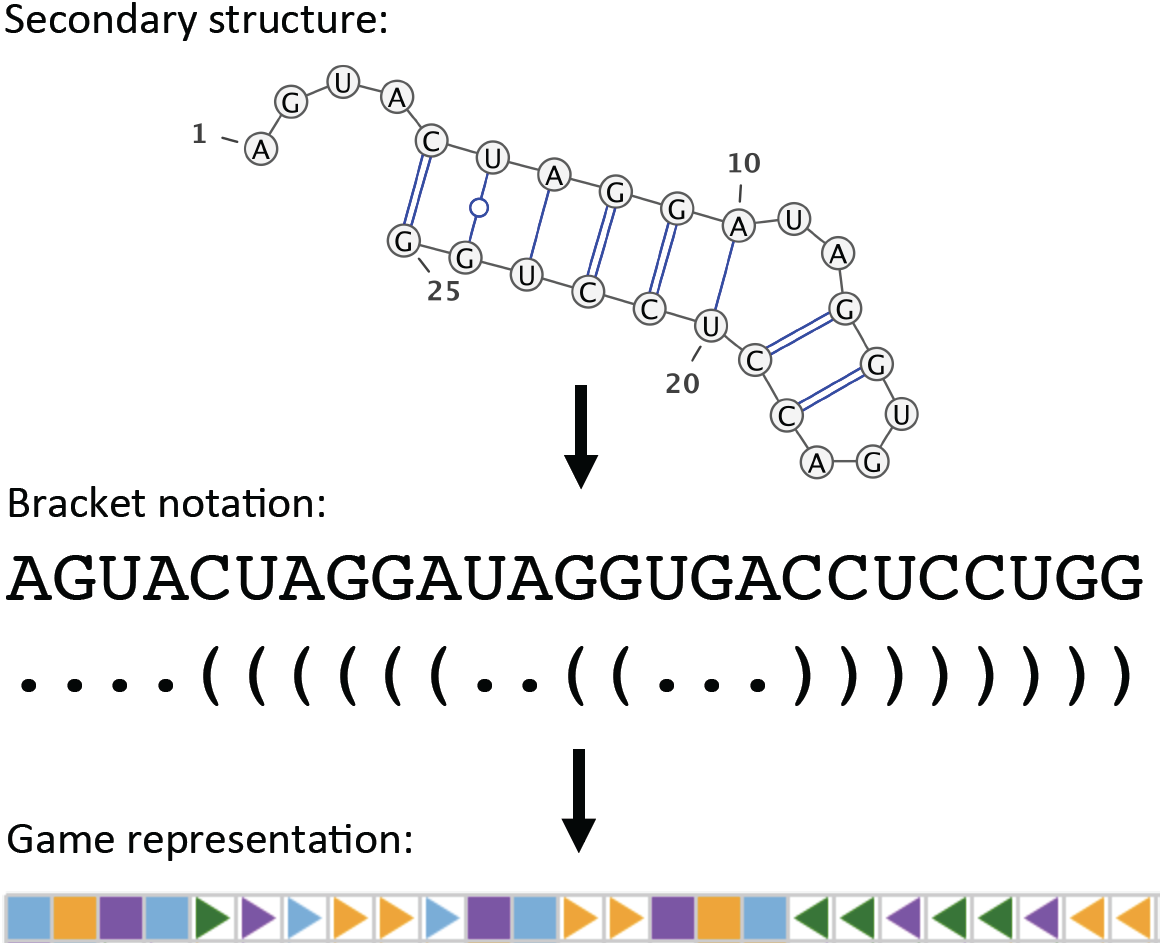
Encoding of RNA sequence and secondary structure in `Ribo`. The secondary structure is drawn with VARNA^46^

### 2.2. Scoring scheme

The scoring scheme that we currently use for evaluating how well the RNA sequences and structures are aligned is based upon the nucleotide sequence scoring scheme derived using a Markovian transition model (States et al. 1991), this approach is how the PAM (Point Accepted Mutation, for proteins) matrices were derived.^35^ The nucleotide scores suggest that for approximately 65% sequence identity, a score between 1.40 and 1.34 should be used for matches and between -1.15 and -1.04 for mismatches. The ratios between these scores convert to approximately the integers +5/-4 for match/mismatch scores. This scoring scheme has been shown to work relatively well for the RNA homology search problem.^36^ We selected relatively low gap-open and gap-extend penalties of -5 and -2 respectively, as indels are thought to be relatively frequent.^37,38^ We add bonuses for matching base-pairs. These bonuses should exceed the penalty for any double mismatches (i.e. *>* 8) resulting from structure-neutral variation (e.g. *A · U* to *G · C*) as well as tolerate the indels that are required to explain base-pair conservation. Therefore a bonus of +12 was selected for aligning base-pairs. At present, covarying sites are not awarded bonuses,^39^ nor are inconsistent and contradicting base-pairs penalized.^40^

### 2.3. Datasets

We used the 5S rRNA multiple sequence alignment (`Rfam` ID RF00001) from the last release of the `Rfam` database^3^ to perform our experiments. This alignment contains 712 sequences and has 231 columns. The 5S rRNA is a component of the large ribosome subunit, and therefore is an essential and ubiquitous RNA, but it has been difficult for `Rfam` to get this alignment correct (data not shown). Although it is a well-known and heavily studied RNA, its alignment remains an open-problem. The 5S rRNA tertiary structure is essential to ribosome assembly and function; hence it is strongly conserved across species. This structure has been experimentally determined as part of the complete ribosome,^41–43^ and this has been used as a reference to obtain the current `Rfam` alignment.

Currently, the grid of the game allows us to represent up to 10 sequences. Thus, we aimed to improve MSA of similar height. We extracted a set of sub-alignments MSA-ref with 4, 6, 8 and 10 sequences. We selected sequences with low average sequence similarity. In our dataset, the average sequence similarity vary from 36% to 58%. This value can be compared with the average sequence similarity of the complete `Rfam` alignment, which is 60%. A full description of the dataset is available in Table 1. This metric is important because sequences with low sequence similarity are hard to align.

**Table 1.** `Ribo` puzzle data set

| Width | Height | ID | Average sequence similarity |  | Percentages in puzzles |  |
| --- | --- | --- | --- | --- | --- | --- |
|  |  |  | Rfam alignment | Ribo puzzle | base pairs | gaps |
| 25 | 4 | 1 | 45 | 56 | 34 | 5 |
| 25 | 4 | 2 | 45 | 33 | 28 | 13 |
| 25 | 6 | 3 | 52 | 73 | 36 | 1 |
| 25 | 6 | 4 | 52 | 58 | 61 | 4 |
| 25 | 8 | 5 | 51 | 61 | 27 | 5 |
| 25 | 8 | 6 | 51 | 54 | 61 | 2 |
| 25 | 10 | 7 | 52 | 68 | 31 | 5 |
| 25 | 10 | 8 | 52 | 47 | 40 | 6 |
| 25 | 4 | 17 | 36 | 33 | 32 | 25 |
| 25 | 4 | 18 | 36 | 40 | 40 | 12 |
| 25 | 6 | 19 | 40 | 47 | 46 | 11 |
| 25 | 8 | 20 | 45 | 56 | 33 | 6 |
| 25 | 10 | 21 | 43 | 48 | 32 | 8 |
| 35 | 4 | 9 | 58 | 73 | 37 | 1 |
| 35 | 4 | 10 | 58 | 65 | 55 | 1 |
| 35 | 6 | 11 | 53 | 59 | 39 | 6 |
| 35 | 6 | 12 | 53 | 51 | 44 | 8 |
| 35 | 8 | 13 | 52 | 67 | 30 | 1 |
| 35 | 8 | 14 | 52 | 44 | 38 | 18 |
| 35 | 10 | 15 | 57 | 68 | 35 | 4 |
| 35 | 10 | 16 | 57 | 59 | 47 | 4 |
| 35 | 4 | 22 | 36 | 34 | 37 | 19 |
| 35 | 4 | 23 | 36 | 40 | 38 | 11 |
| 35 | 6 | 24 | 40 | 34 | 32 | 22 |
| 35 | 6 | 25 | 40 | 43 | 40 | 11 |
| 35 | 8 | 26 | 45 | 51 | 34 | 6 |
| 35 | 10 | 27 | 43 | 49 | 38 | 8 |

We built the test sets with the sequences from the original `Rfam` alignment not used in the reference sub-alignment set MSA-ref. Hence, the discriminative power of a sequence of 6 sequences has been estimated on a benchmark set containing the other 706 sequences from the `Rfam` alignment.

### 2.4. Puzzle construction

We built two sets of puzzles with 25 and 35 columns, labeled as “Easy” and “Hard”, from the `Rfam` sub-alignments MSA-ref described above. The choice of the sizes has been determined to maximize the use of the grid of 50 columns currently used in our game, and at the same time to give the players enough room to explore the configuration space.

Because the experimentally determined structure may not always be available, we ignored the consensus structure that is available in the original `Rfam` alignment. Instead, we predicted a secondary structure using the maximum expected accuracy (MEA) secondary structure predicted by `RNAfold`^44,45^ for each individual sequence. Therefore, it is important to keep in mind that in this study, the `Rfam` alignments benefit of an information not used by the players. We removed empty columns (i.e. columns containing only gaps) from the sub-alignments, and extracted all continuous regions of 25 and 35 columns. Then, we removed from each region the base-pairs that were not included within this region. In other words, if a nucleotide has a predicted interaction with another nucleotide outside of the region of interest, this base-pair is ignored.

We sorted all regions according to the total number of base-pairs in the region (i.e. the sum of the number of valid base-pairs predicted by `RNAfold` within the region for each sequence of the sub-alignment). Regions without any base-pair were ignored. Finally, we selected first the region with the larger number of base-pairs, then the next one provided that it does not overlap with the region previously selected, until the queue is left empty. In the end, we generated 27 puzzles that are described in Table 1. In this table, we report the number of columns (width) and sequences (height) of the puzzle, and its ID in the game. We also report the average sequence similarity of the `Rfam` sub-alignment used to create the puzzle, as well as the average sequence similarity of the puzzle. Finally, we report the percentage of nucleotides involved in a base pairing interaction, and the percentage of gaps found in the initial configuration of the puzzle.

### 2.5. Benchmarking methodology

The quality of the alignments was evaluated using `Infernal`.^31^ `Infernal` is the software suite used to build the covariance models from `Rfam` seed alignments and search for homologs (available in the full `Rfam` alignment).

For each submission (i.e. a puzzle solved by a player), we substituted the original alignment of the region used to build the puzzle, with the solution provided by the player. Then, we calculated a covariance model for each of these alignments (the original one and the one built using the submission) with the program `cmbuild` of the `Infernal` package. Finally, we calibrated the covariance models with the program `cmcalibrate`, and used the program `cmsearch` to compute a fitness score evaluating the likelihood of the covariance model on each sequence of the test set (the set of the `Rfam` seed sequences not used in the original alignment).

The fitness is estimated with the E-value calculated by `cmsearch`. In our experiments, we report the average E-value of all sequences in the test set. Among all solutions collected for a given sub-alignment, only the best values are reported. Indeed, as in `Phylo` the purpose of our system is to generate a sparse set of potential solutions in which we have high probability to find a configuration improving the original one.

## 3. Results

### 3.1. Game design

`Ribo` is inspired from our previous contribution `Phylo`. We abstract a sequence alignment into a tile-matching game, where nucleotides are represented with coloured bricks that can be moved horizontally on a grid. The objective of the players is to align the nucleotides of similar colours within the same columns, in order to reveal similarities between sequences.

Nonetheless, RNA alignments have a major difference with DNA alignments. The conservation of the native (functional) structure is often more important than the conservation of the primary sequence. As in `Rfam`, the molecular structure are represented by secondary structures. Hence, RNA alignments aim to conserve base-pairs. For instance, if a base-pair occurs between indices (*i*_1_*, j*_1_) in one sequence and another one between indices (*i*_2_*, j*_2_) in a second sequence, then the alignment of nucleotides at index *i*_1_ and *i*_2_ in the same column must be, as much as possible, associated with the alignment of nucleotides at index *j*_1_ and *j*_2_. Since the conservation of base-pairing properties is essential for RNA alignments, we need to design new mechanisms to represent this information and enable users to use it in the game.

RNA secondary structures encompass the maximal set of stem and stem-loops formed by canonical base-pairing interactions (Watson-Crick and Wobble). Each nucleotide can be involved in at most one base-pair (i.e. no base-triples) and crossing interactions are forbidden (i.e. no pseudo-knots).

In `Ribo`, we chose to adapt the bracket notation frequently used to represent RNA secondary structures. An open parenthesis indicates that the nucleotide is paired with the first available nucleotide associated with a close parenthesis on its right. Dots represent unpaired nucleotides. The bricks used in `Ribo` merge the sequence and structural information into a single token. As in `Phylo`, the colour of the brick encodes the type of the nucleotide (i.e. A, C, G or U). In addition, we use now a new set of bricks with different shapes to encode the base-pairing properties. Hence, a triangle pointing to the right indicates that the nucleotide is paired with another nucleotide on its right (i.e. the equivalent of the open parenthesis), while a triangle pointing to the left indicates the opposite. Unpaired nucleotides are represented using a squared brick. Figure 1 illustrates our encoding.

The scoring scheme used in `Ribo` significantly diverges from the one used in `Phylo`. Since the phylogenetic tree is unknown (as it is often the case in RNA alignments), we use a sum-of-pair scoring function. In other words, the total score of a multiple sequence alignment is the sum of the alignment scores of each pair of sequences. Here, matches (bricks with identical colours aligned together) receive a bonus of +5, and mismatches (bricks with different colours) receive a penalty of -4. The opening of a gap costs -5 and their extension only -2. Finally, the alignment of a base-pair receive a bonus a +12. We motivate these choices in section 2.2.

As it could be the case in practical bioinformatics applications, in `Ribo` we do not penalize misaligned or contradicting alignments of base-pairs.^47^ We argue that including such penalties would affect the design of the game and diminish the engagement of players. By contrast, the bonuses assigned to matched base-pairs create an extra incentive to players to explore the configuration space, and is sufficient to serve our purpose to align secondary structures.

We show a screenshot of the game in Figure 2. The game board uses some successful element designs previously developed with `Phylo` such as the score bar of top indicating the current score, best score achieved during this session and score to beat (i.e. the Par). We also added new features such as the locks on the left side. The latter enables the players to “lock” a row and move the full sequence as whole, preserving gaps between the bricks. This feature aims to facilitate the playability of the game on tablets and mobile devices. Moreover, the visual identification of long-range base-pairs can be difficult. To address this issue, we implemented a highlight mechanism that shows the base-paired brick every time that the cursor overlaps with a brick. Finally, we do not need to represent a phylogenetic tree as it was the case in `Phylo`. Thus, we have more space to display the game board. With `Ribo`, we decided to increase the grid to 50 columns (instead of 25 with `Phylo`). This upgrade is essential because base-pairs can involve nucleotides that are very distant in the sequence. `Ribo` is available at http://ribo.cs.mcgill.ca.

**Fig. 2.**
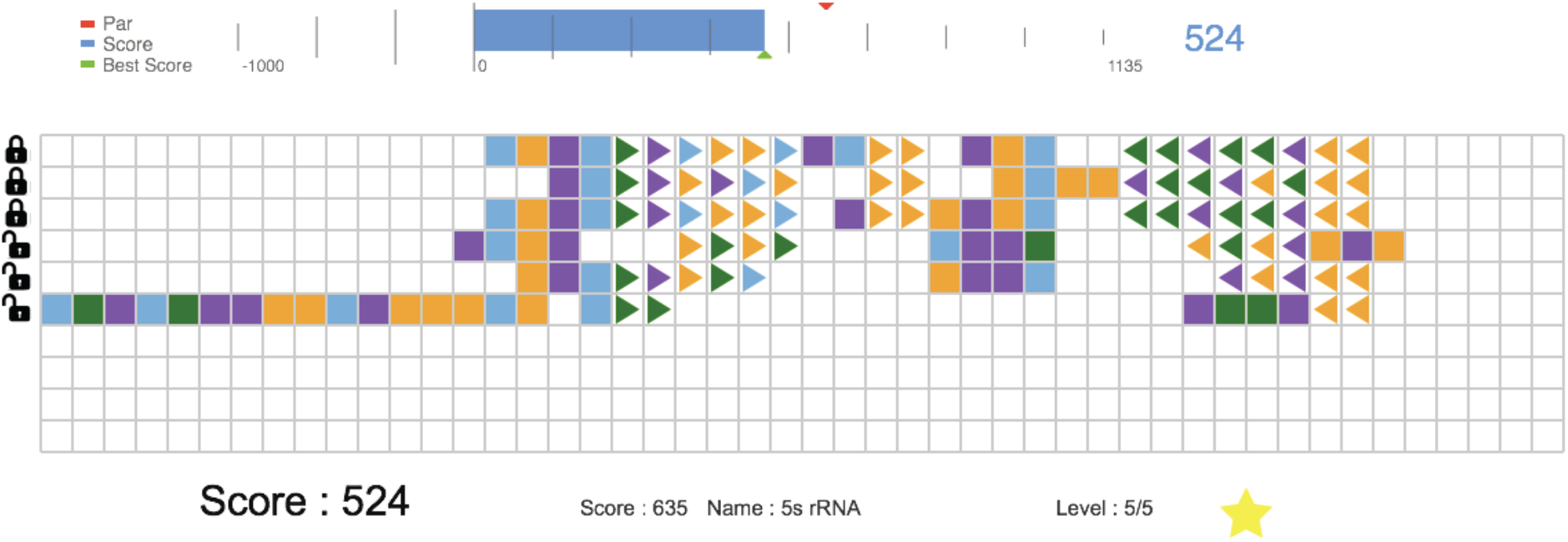
Screenshot of `Ribo`

The progression of the player within the game is similar to the one used in `Phylo`. First, the user starts with two sequences and tries to find an alignment with a score that is at least as good as the one found in the original alignment. Once this milestone is reached, the player can access the next stage and add one more sequence to the game. The game is complete when all sequences have been added and when the player managed to beat or match the score of the original alignment.

### 3.2. Game statistics

Approximately 15 players recruited from undergraduate and graduate students in computational biology at McGill and University of Canterbury participated to the study. We collected 115 submissions (i.e. puzzles completed) whose distribution is detailed in Table 2. The “easiest” puzzles (least numbers of rows and columns) have been significantly more played than the others. It had to be expected since all participants were beginners and thus needed to learn the rules and train on easy instances first. Nonetheless, it is worth noting that the majority of participants was not familiar with RNA alignments before starting to play.

**Table 2.**
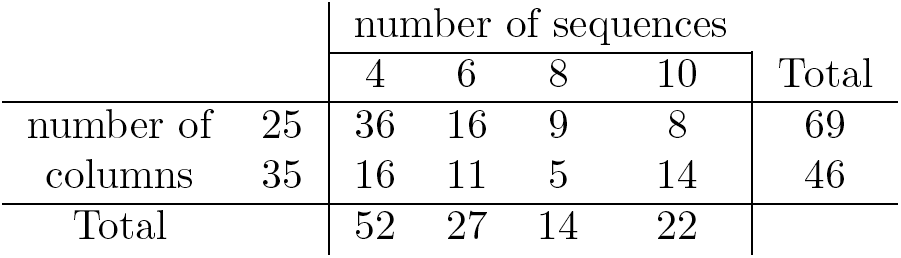
Number of solutions collected

|  |  | number of sequences |  |  |  | Total |
| --- | --- | --- | --- | --- | --- | --- |
|  |  | 4 | 6 | 8 | 10 |  |
| number of | 25 | 36 | 16 | 9 | 8 | 69 |
| columns | 35 | 16 | 11 | 5 | 14 | 46 |
| Total |  | 52 | 27 | 14 | 22 |  |

### 3.3. Performance

In Figure 3(a), we report the average E-values obtained on puzzles with the same number of sequences. The decrease of E-values observed on the alignments improved by the gamers tends to validate our approach. An exception is for alignments with 6 sequences. This discrepancy is most likely due to incorrectly predicted base-pairs that resulted in alignment of worse quality. We also note a trend toward higher average E-values when the number of sequence increases. This phenomenon could be an artifact of the small sample set, but could also reflect a real phenomenon. Since all the sequences are quite diverse, higher numbers of sequences in the alignments result in lower the probabilities for each state in the covariance model. Consequently the E-values are likely to be higher.

**Fig. 3.**
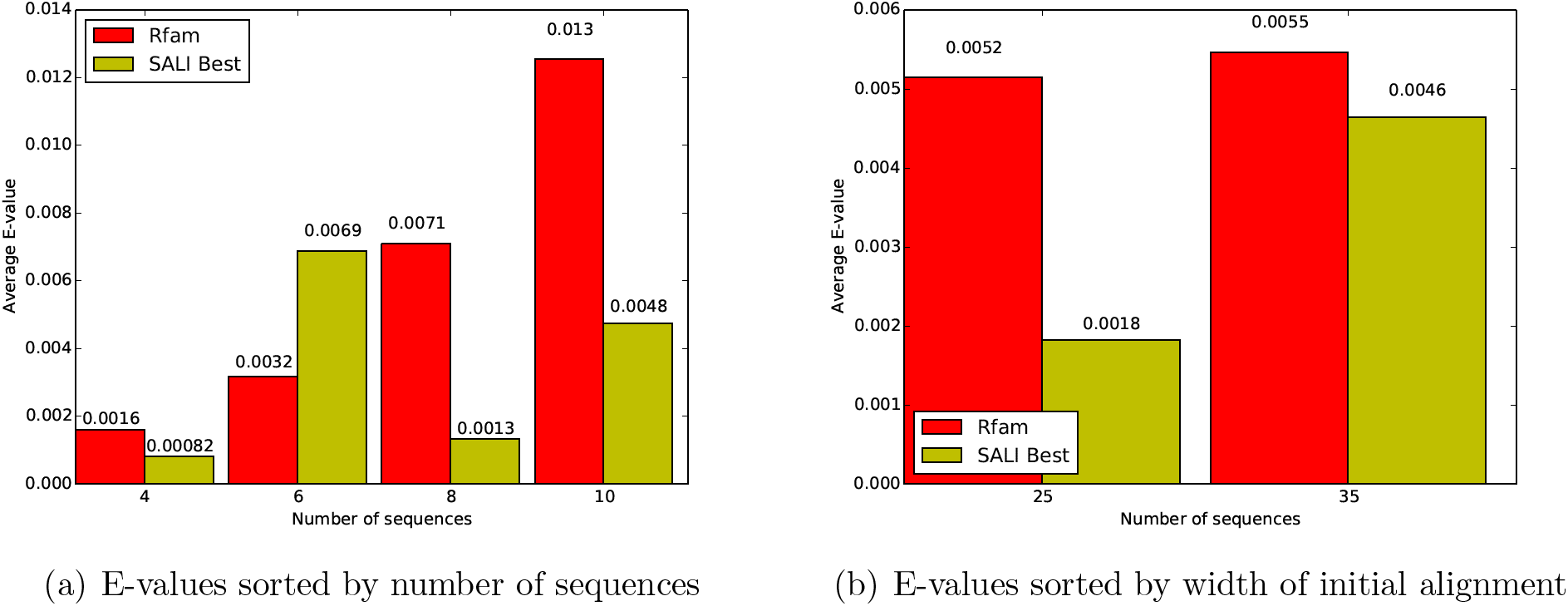
Average E-values of the covariance model on sequences in the test set. Average E-values calculated with the covariance model obtained from the `Rfam` alignments are shown in red, and average E-values calculated with the covariance model obtained from `Ribo` alignments are displayed in yellow. The left panel shows the results obtained when the puzzles are sorted by number of sequences, while the right panel shows the same data when the puzzles are sorted by number of columns (i.e. defined as difficulty in the game).

Similar observations can be done when the puzzles are sorted by difficulty in Figure 3(b) (i.e. the number of columns of the regions used to build in them). The average E-values decrease for the Easy (25 columns) and Hard puzzles (35 columns). However, the magnitude of the improvement is less pronounced for the largest puzzles. A lower number of submissions (See Table 2) as well as higher difficulties to solve these puzzles can justify this difference.

The data presented above suggest that overall our game has the potential to improve RNA alignments. Nonetheless, the distribution of E-values also needs to be considered to understand the real performance of this methodology. Indeed, our data show that E-values obtained with the covariance models calculated from the alignments generated by gamers were better (i.e. lower) on lowest ranked sequences (i.e. sequences with the worst fit to the covariance model), than the E-values obtained on the same sequences with the covariance model calculated from the original alignment. By contrast, the E-values obtained with the covariance model obtained from the original alignment are better than those obtained with the alignment improved by the game on highest ranked sequences (i.e. the sequences with the best fit to the covariance model). Therefore, the new covariance models appear to outperform the ones built from `Rfam` sub-alignments for recognizing distant homologs, but may lack the specificity of the latter to identify sub-families.

## 4. Discussion

The accurate alignment of sequences for structural RNAs remains a challenging problem. The “gold standard” remains the manual construction of alignments. In fact, the accuracy of careful manual comparisons of sequences were shown to be 100% accurate when evaluated against structures derived from crystallographic data.^32^ However, this approach is very time consuming and requires highly trained and committed individuals.

We have shown the potential for “crowd sourcing” the RNA multiple sequence alignment problem. Alignments can be broken into a series of sub-sequences and sub-alignments. Crowd-sourced solutions to these can be stitched together, thus building up reasonable solutions to computationally challenging problems. This paper is a proof-of-concept that crowd-sourcing techniques can be used to maintain and improve public RNA alignment repositories.

Feedbacks from our players collected after the benchmark suggest that the limit of the human-computing system have not been reached yet. In particular, we could increase the number of sequences and columns. Nonetheless, such upgrades will also require the development of new GUI features to help the player to deal with large data sets, and help them to efficiently explore the conformational space. For instance, advanced visualization tools to display long-range base-pairing interactions.

We argue that more sophisticated strategies to build the puzzles have the potential to increase the performance of our crowd-computing system. Indeed, the puzzles used in this study are built from continuous regions of an `Rfam` alignment with 35 columns. This strategy prevents us to use long-range interactions between nucleotides that are separated by more that 35 positions. This is an important issue if we wish to use `Ribo` to align multi-loop regions of RNA with sophisticated secondary structures. To address this problem, we suggest building puzzles from discontinuous regions of a full alignment. For instance, we can concatenate a region with 20 columns with another region of 20 columns that contains the nucleotides predicted to base-pair with those of the first region.

Although relatively rare, pseudo-knotted can carry important functions. Currently, 89 `Rfam` families among 2208 have pseudo-knot annotations. To handle these families, a second set of parenthesis could be used to represent interleaved interactions.

Finally, due to the broad interest of the scientific community in obtaining accurate RNA alignments, we can envision the use of `Ribo` as a web widget on research web sites to promote the understanding on RNA research to a broad public and engage citizen scientists. The deployment of an open crowd-computing platform such as `Open-Phylo`^48^ is also scheduled.

## 5. Availability

`Ribo` can be played at http://ribo.cs.mcgill.ca. The source code and data used in the project are also freely accessible at http://jwgitlab.cs.mcgill.ca/arthurkam/rna-phylo.

